# Chronic pain mediated changes in the appetitive value of affective gentle touch in mice

**DOI:** 10.1101/2023.11.09.566431

**Authors:** Maham Zain, Laura Bennett, Hantao Zhang, Quinn Pauli, Juliet Arsenault, Shajenth Premachandran, Jenny Cheung, Christine Pham, Lexi Wowk, Sékou-Oumar Kaba, Samuel Fung, Irene Lecker, Eric Salter, Feng Wang, Reza Sharif-Naeini, Yves De Koninck, Robert P. Bonin

**Affiliations:** University of Toronto; McGill University; University of Laval; CERVO Brain Research Centre

## Abstract

The existence of skin-to-brain circuits for rewarding gentle touch highlight its critical nature across species. However, this gentle affective touch is not always appetitive and can produce aversion or negative affect in disorders such as chronic pain. Sensory neurons expressing the protein MrgprB4 detect gentle stroking in mice and their activation of these neurons known to be positively reinforcing. Here we assess whether activation of channelrhodopsin (ChR2) expressing MrgprB4 afferents signal positively valenced tactile information and whether this is altered in models of chronic pain. We further interrogate how this this sensory information is reflected in the downstream circuits recruited. Optogenetic activation of MrgprB4 lineage afferents was found to be appetitive in control and capsaicin sensitized mice but not nerve injured mice, indicating that the appetitive value is diminished in neuropathic pain. Remarkably, this appetitive value was partially recovered in male nerve injured mice by treatment with the analgesic gabapentin. These behavioral changes were also accompanied by different patterns of neuronal activity throughout the brain, including altered activation of sites that receive direct projections from the spinal cord between sham and nerve injured mice. Together, these findings highlight the plastic nature of these affective tactile circuits under pathological conditions.

## Introduction

The somatic sensations of touch and pain are complex multi-dimensional sensory modalities. The sensory discriminative dimension enables us to differentiate and identify the spatial and temporal characteristics of a stimulus. The affective-motivational dimension attaches emotional or hedonic value to sensory stimuli and can often guide behavior in aversive and appetitive ways. These homeostatic drives, with respect to tactile sensations, are critical for survival as they ensure the promotion of behaviours that allow us to escape harmful stimuli and gravitate towards evolutionarily beneficial stimuli.^1–4^ This includes aversive motivation to escape pain and appetitive motivation towards pleasant touch.^4–6^

Despite this polarity in motivational valence the distinction between aversive pain and appetitive touch can become blurred in various pathological conditions. In fact, the relationship between pain and pleasant touch is complex and these sensations can modulate one another. While gentle touch has the potential to be analgesic^7–9^, the presence of pain pathologies may render it aversive, or at the minimum not pleasant^8,10,11^. A prominent example is in chronic pain disorders like fibromyalgia, where patients develop anhedonia to typically pleasant gentle touch^12,13^. This loss of gentle touch reward contributes heavily to the disease burden and significantly affects patient quality of life^14^. While there has been recent work identifying mechanisms that drive touch reward, there is limited understanding as to how this is altered in pain states and mechanisms underlying this plasticity^15^.

Coding of emotionally valenced sensory information begins at the periphery such that signals transmitted by specific sub-classes of primary afferents give rise to sensory perceptions with motivational value.^10,16,17^ Classic microneurography studies in humans had identified C-tactile afferents as key players in the transmission of pleasant slow stroking touch.^10,18–20^ A comparative class of low threshold C-fibers has also been identified in rodents. These afferents can be distinguished through their expression of a G-protein coupled receptor, MrgprB4. Activation of these afferents in mice is positively reinforcing.^16,21^ While recent work has implicated this group in physiologically relevant behaviors^15^, little is known about the plasticity of the behavioral phenotype associated with the activation of MrgprB4-lineage afferents. This further begs the question as to whether the behavioral response is at least partially mediated by the differential recruitment of downstream targets, both at the spinal and supraspinal level. Using optogenetics, behaviour, immunohistochemistry, and electrophysiology we sought to examine the robustness of the appetitive response associated with activation of MrgprB4-lineage afferents in various pain models, and elucidate CNS neuronal activity associated with these responses.

## Results

### Characterizing expression of ChR2 in the MrgprB4-ChR2 transgenic mouse model

To gain optogenetic control of MrgprB4 lineage neurons, we crossed micecontaining the Cre-recombinase gene under the control of a MrgprB4 promoter (MrgprB4-Cre), with mice expressing a loxP-flanked STOP cassette upstream of a ChR2-EYFP fusion gene at the Rosa 26 locus (Rosa-CAG-LSL-ChR2(H134R)-EYFP-WPRE; stock number 012569, The Jackson Laboratory, Bar Harbor, ME). The resulting mice express Channelrhodopsin (ChR2) in neurons that expressed MrgprB4 at any point in development (MrgprB4-ChR2). The presence of ChR2 was confirmed through immunostaining for YFP, where YFP+ afferents were observed to innervate lamina II of the superficial dorsal horn in the spinal cord in a manner consistent with non-peptidergic afferents positive for Isolectin B4 (IB4) (SI Appendix, **Fig. S1A**). Quantitatively, more than half of the ChR2 labelled population were also positive for IB4 (SI Appendix**, Fig. S1B and C**). We also observed some overlap between the ChR2 and CGRP positive peptidergic dorsal root ganglions (DRG) given that MrgprB4 is more broadly expressed during the first few postnatal days than in adults (SI Appendix**, Fig. S1D and E**). These results indicate successful labeling of MrgprB4 lineage afferents and are consistent with other reports on this mouse line^15,22^.

### MrgprB4 lineage afferent mediated affective touch signalling does not interact with acute somatosensation or acute pain

Given the challenges associated with concurrent peripheral light delivery and sensory testing, we initially employed the use of an epidural optic fiber to deliver light to the central terminals of the primary afferents originating from the hindlimbs^23^. Activation of lumbar MrgprB4 afferents by epidural delivery of blue light did not change thermal or mechanical thresholds in the hind paws of implanted mice (**Fig. 1A and B**). Additionally, intraplantar injection of capsaicin did not affect acute observable behavioral responses to peripheral light stimulation of the hairy skin of the injected paw (**Fig. 1C**) despite capsaicin induced mechanical allodynia as indicated by von Frey (**Fig. 1D**). This suggested that sensitization resulting from insult to an area with no MrgprB4 afferent arborizations did not modify acute responses to activation of MrgprB4 lineage afferents.

**Figure 1.**
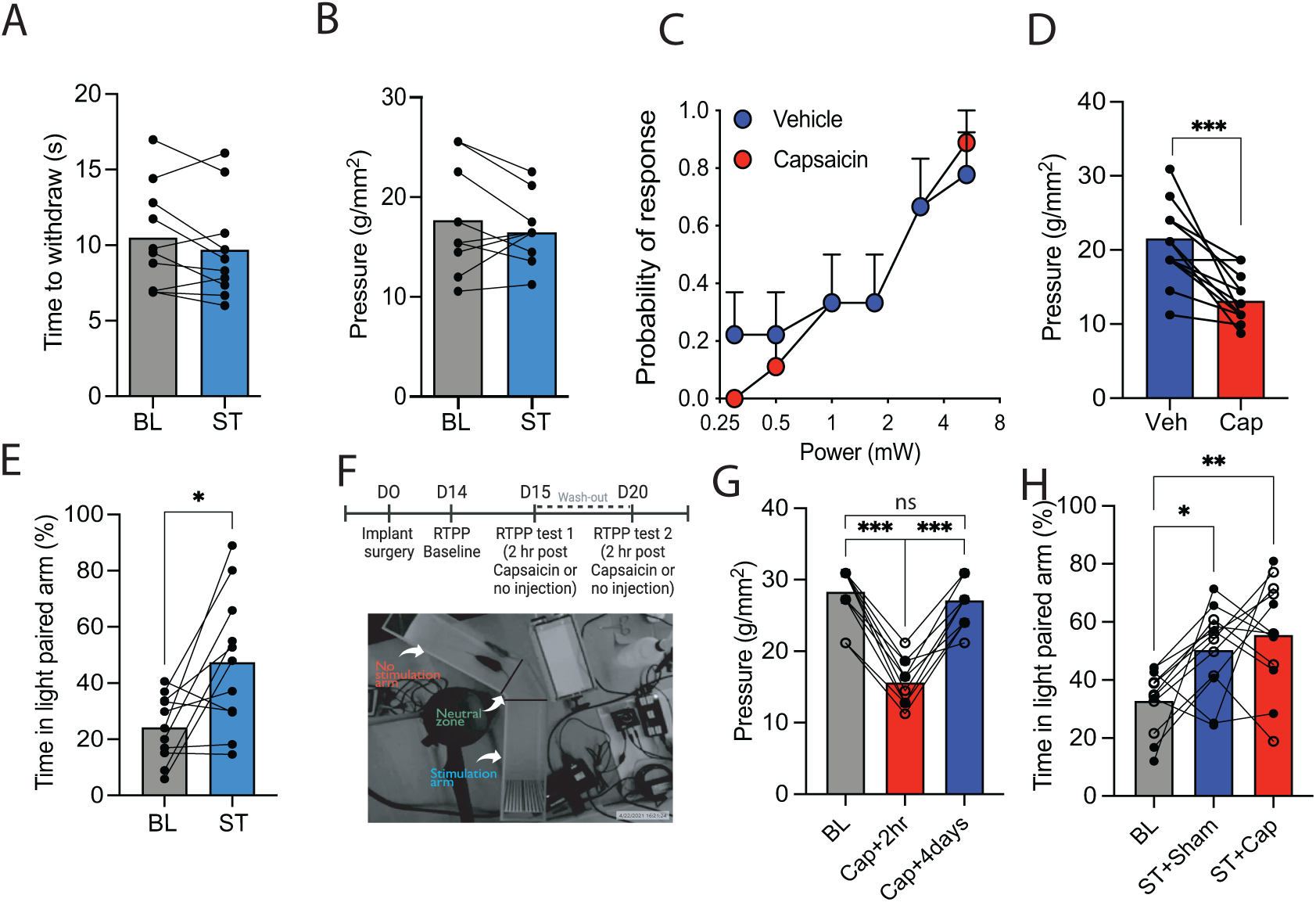
MrgprB4 lineage afferent mediated affective touch signalling does not interact with acute somatosensation or acute pain. **(A-B)** Thermal **(A)** and mechanical thresholds **(B)** unchanged before and during stimulation of MrgprB4 lineage afferents through an epidural implant. Paired t-test (p > 0.05; n = 10). **(C)** Acute behavioral responses to peripheral blue light unchanged in MrgprB4-ChR2 mice that received either a vehicle or Capsaicin injection. Log-rank test (p > 0.05; n = 9). **(D)** This capsaicin injection induced mechanical allodynia. Unpaired t-test (***p < 0.001; n = 13) **(E)** Stimulation of the MrgprB4 lineage afferents through a spinal implant in a RTPP assay could increase preference for the light paired arm compared to baseline when no stimulation was provided. Paired t-test (*p < 0.05; n =11). **(E)** Experimental timeline (cross-over design) to test capsaicin injected MrgprB4-ChR2 mice in RTPP and pictorial representation of the setup used. **(G)** The capsaicin injection produced mechanical allodynia that resolved in the number of days that mice were given between repeated RTPP trials. RM one-way ANOVA with Sidak multiple comparison test (*F*_(1.8,20.1)_ = 84.12, \*\*\**p* < 0.001; BL vs Cap+2hr ***p < 0.001, Cap+2hr vs Cap+4days ***p < 0.001, BL vs Cap+4days p > 0.05, n = 12). **(H)** An increase in preference for the light paired arm was observed in response to stimulation irrespective of sex and whether the mice received a sham procedure that did not include an injection or the capsaicin injection. RM two-way ANOVA with Sidak multiple comparison test (effect of treatment: F_(2, 20)_ = 6.748, **p < 0.01; effect of sex: F_(1, 10)_ = 0.3, p > 0.05; interaction effect: F_(2, 20)_ = 0.09, p > 0.05; BL vs ST+Sham *p < 0.05, BL vs ST+Cap **p < 0.01; n = 12). Open circles = females; Closed circles = male BL, Baseline; ST, Stimulation; Cap, Capsaicin; ANOVA, Analysis of variance; RM, Repeated measures; RTPP, Real time place preference.

To assess the appetitive quality associated with activation of MrgprB4 lineage afferents we performed a variation of a blue light paired real time place preference (RTPP) assay. We used a spinal implant to deliver light to the central projections of the primary afferents that originate from the hindlimb, a strategy that can reliably produce optogenetic responses without any motor or sensory deficits.^24^ Preference for the initially non preferred (INP) arm could be increased by delivering blue light stimulation in that arm in MrgprB4-ChR2 male mice (**Fig. 1E**). To assess whether this appetitive value was maintained in females and whether it was sensitive to acute pain induced via capsaicin we performed real time place preference in implanted male and female mice that had received an intraplantar capsaicin injection. Mice were tested in a counter balanced cross over study where they were either given no injection (sham) or a capsaicin intraplantar injection with a 4-day washout period (**Fig. 1F**). The dose of capsaicin used was found to reliably elicit mechanical allodynia, as measured by von Frey, that resolved in 4 days (**Fig. 1G**). Despite this sensitization, preference for the INP could still be increased when the mice were injected with capsaicin, like the increase observed when they received no injection (**Fig. 1H**). Sex was not a significant variable in the experiment indicating that this was not a sexually dimorphic response.

### Appetitive responses to MrgprB4 lineage afferent activation are modulated by nerve injury

Remarkably, unlike acute injury via capsaicin, we observed that the appetitive value of activation of the MrgprB4 afferents was sensitive to nerve injury induced via a spared nerve injury (SNI). We performed RTPP in implanted male mice that had undergone an SNI or sham surgery 7 days prior (**Fig. 2A**). SNI is known to reliably induce peak mechanical hypersensitivity that emerges as early as 2 days post-surgery and is maintained for several months^25–27^. Sham surgery mice received all surgical manipulation except for the ligation and transection of the appropriate sciatic nerves. In contrast to their sham counterparts, the SNI animals did not show an increase in preference in response to light stimulation in the INP arm of the assay (**Fig. 2C**). Additionally, the SNI animals, in contrast to sham mice, had lower number of entries into the light paired arm during stimulation compared to baseline (**Fig. 2D**).

**Figure 2.**
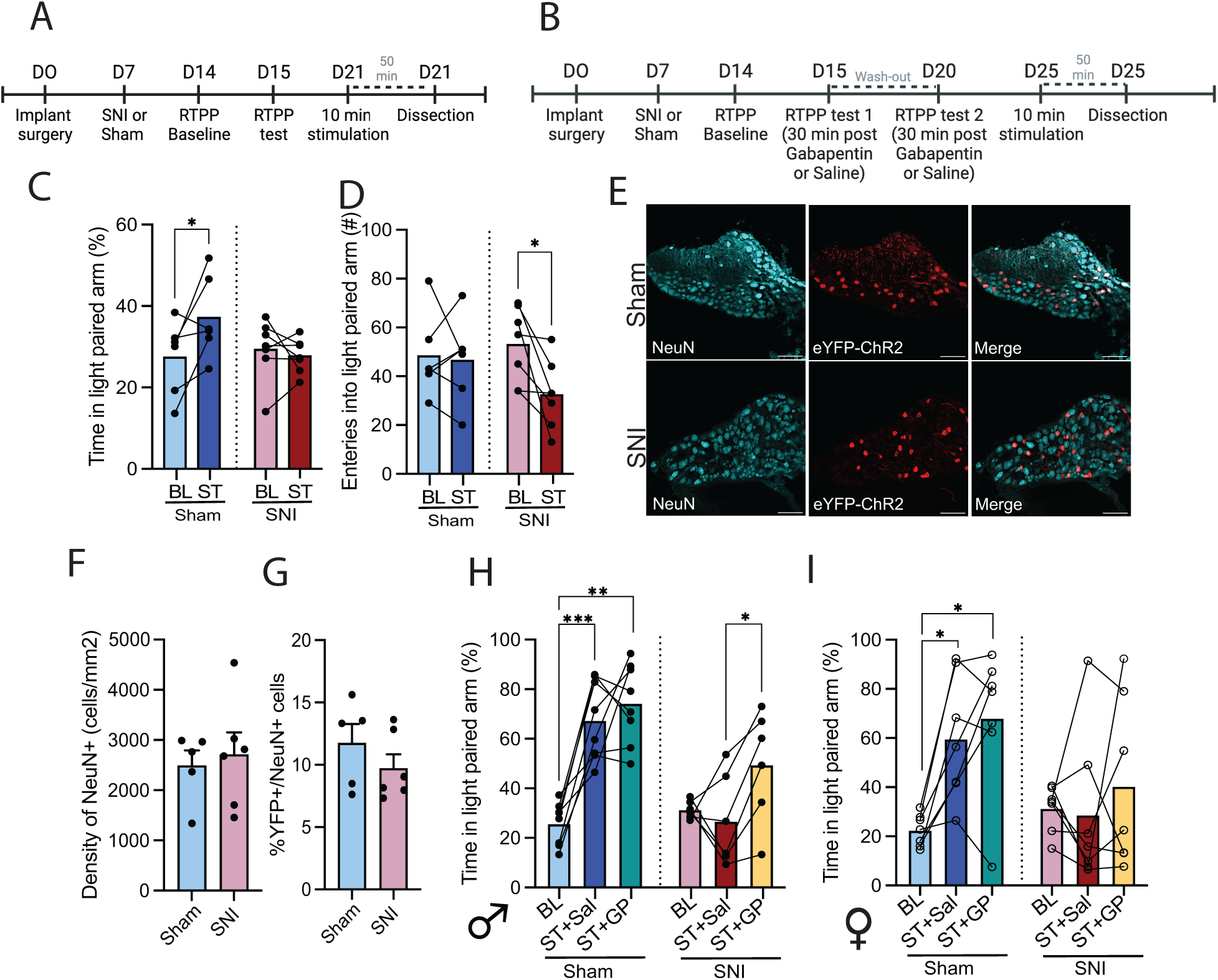
Neuropathic pain induced plasticity in the appetitive value of affective touch mediated by MrgprB4 lineage afferents. **(A)** Experimental timeline to study the effect of nerve injury on blue light preference in MrgprB4-ChR2 **(B)** Experimental timeline (counter balanced cross over study design) to test the effects of gabapentin on blue light preference in nerve injured and sham MrgprB4-ChR2. (**C**) An increase in preference for the light paired arm was observed in response to stimulation in the sham mice but not the SNI mice. RM two-way ANOVA with Sidak multiple comparison test (effect of trial: *F*_(1, 11)_ = 2.9, p > 0.1; effect of surgery: *F*_(1, 11)_ = 1.1, p > 0.1; interaction effect: *F*_(1, 11)_ = 5.8, p < 0.05; Sham-BL vs Sham-ST *p < 0.05; n = 6-7 mice per gp) (**D**) A decrease in number of entries into the light paired arm were observed in the SNI mice during the stimulation trial compared to baseline and this decrease was not observed for the sham mice. RM two-way ANOVA with Sidak multiple comparison test (effect of trial: *F*_(1, 11)_ = 5.2, p < 0.05; effect of surgery: *F*_(1, 11)_ = 0.40, p > 0.5; interaction effect: *F*_(1, 11)_ = 3.7, p > 0.05; SNI-BL vs SNI-ST *p < 0.05; n = 6-7 mice per gp). (**E**) Representative images showing NeuN and eYFP-ChR2 expression in lumbar DRGs. (**F, G**) No differences in total density of neurons (**F**) or percent of YFP expressing neurons **(G)** in lumbar DRGs of SNI and sham MrgprB4-ChR2 mice. Unpaired t-test (p > 0.05; n = 5-6 mice per group, 1-2 slices per mouse). (**H**) An increase in preference for the light paired arm was observed in response to stimulation in the sham male mice but not the SNI male mice; however, administration of gabapentin in SNI male mice resulted in a higher preference compared to when the mice were injected with saline. RM two-way ANOVA with Sidak multiple comparison test (effect of trial: *F*_(1.9, 22.3)_ = 19.2, p < 0.0001; effect of surgery: *F*_(1, 12)_ = 13.5, p < 0.01; interaction effect: *F*_(2, 24)_ = 9.5, p < 0.001; Sham-BL vs Sham-ST+Sal ***p < 0.001, Sham-BL vs Sham-ST+GP **p < 0.01, SNI-ST+Sal vs. SNI-ST+GP *p < 0.05; n = 6-8 mice per group) (**I**) An increase in preference for the light paired arm was observed in response to stimulation in the sham female mice but not the SNI female mice and these effects were maintained regardless of drug administration. RM two-way ANOVA with Sidak multiple comparison test (effect of trial: *F*_(1.9, 22.9)_ = 6.2, p < 0.01; effect of surgery: *F*_(1, 12)_ = 2.8, p > 0.05; interaction effect: *F*_(2, 24)_ = 3.9, p < 0.05; Sham-BL vs. Sham-ST+Sal *p < 0.05, Sham-BL vs. Sham-ST+GP *p < 0.05; n = 7 mice per group) Open circles = females; Closed circles = male; GP, Gabapentin

These differences existed despite no difference in locomotion between the experimental groups in both the baseline and the stimulation trial (SI Appendix**, Fig. S2**). Effects of the SNI surgery were also confirmed through immunostaining of ATF3 (SI Appendix**, Fig. S2B and D**), a nerve injury marker whose expression is known to correlate with mechanical hypersensitivity. A limited fraction (<15%) of the Channelrhodopsin labelled population expressed ATF3, with no difference between the 2 surgical groups (SI Appendix**, Fig. S2C and D**). Furthermore, to confirm that the loss of light preference in our SNI mice was not driven by a loss in the number of Channelrhodopsin expressing neurons we quantified the density of neurons and the percentage of ChR2 expressing neurons in lumbar DRG sections from both groups of animals. Consistent with recent reports^28^ we did not observe a decrease in the density of neurons (**Fig. 2E and F**) nor the percentage of ChR2 expressing neurons (**Fig. 2E and G**) in the mice that had undergone SNI compared to sham surgeries.

### Partial recovery of appetitive value with gabapentin in nerve injured male mice

No loss of the MrgprB4 lineage afferents hinted to the possibility of recovering the appetitive value of affective touch in nerve injured animals. Gabapentin is a commonly prescribed analgesic for neuropathic pain in humans and has been shown to be analgesic in spared nerve injury models in mice in our lab, in addition to many others. To this effect, we repeated the previous study in a new cohort of male and female implanted mice. In this counterbalanced crossover study, the SNI and sham animals received saline and gabapentin with a 4-day wash out period (**Fig. 2B**). Consistent with previous results we once again found that sham male mice had increased preference for the light paired arm in response to stimulation, compared to baseline, regardless of the drug administered. However, male SNI mice given gabapentin exhibited a higher preference for the light paired arm during stimulation, compared to saline-injected male SNI mice (**Fig. 2H**). We also found a loss of appetitive affective touch in female SNI mice, though this effect was not modulated by the administration of gabapentin (**Fig. 2I**). This indicates at least partial recovery of the appetitive value of touch in gabapentin-treated male, but not female, mice with a nerve injury.

### Appetitive responses to MrgprB4 lineage afferent activation are modulated by disinhibition in spinal cord circuits

The central sensitization caused by disinhibition of spinal cord circuits is a key substrate underlying abnormal touch processing in neuropathic pain. This disinhibition is a byproduct of the downregulation of the K+-Cl− co-transporter KCC2 that results in a depolarizing shift in the Cl− reversal potential thus impairing GABAA/GlyR-mediated inhibition in a sex independent manner^29^. To disentangle the mechanistic basis of the loss of the appetitive value of touch in nerve injured mice we performed a variation of the real time place preference assay in MrgprB4-ChR2 mice that were given an intrathecal injection of DIOA, a KCC2 blocker, to induce pharmacological disinhibition. Given the challenges of performing an intrathecal injection in mice with spinal implants, we switched to a peripheral method of light delivery for optogenetic activation of our afferents (SI Appendix**, Fig. S3A**). We created a rectangular chamber with a LED-illuminated floor where half of the floor was illuminated with orange light and the other half with blue light. We found that mice injected with DIOA preferred to spend less time in the blue side of the assay when the LEDs were active compared to baseline, when the LEDs were switched off (SI Appendix**, Fig. S3B**). Such a decrease was not observed when the mice were injected with saline, though we did not observe an increase in preference either as would be expected with the activation of MrgprB4 lineage afferents (SI Appendix**, Fig. S3B**). This may have been, in part, driven by the preferential aversion mice display to blue peripheral light, which we observed in this experiment and has also been reported by others using similar paradigms^30^.

### Characterization of the spinal dorsal horn neurons recruited following activation of MrgprB4 lineage afferents in nerve injured and control mice

Collectively our findings have highlighted that disinhibition, even when restricted to the spinal level, is sufficient to alter the appetitive value of affective touch that is mediated by MrgprB4 lineage afferents. To characterize the spinal dorsal horn neurons that the MrgprB4 lineage afferents provide input into at baseline conditions, we conducted whole cell recordings of superficial dorsal horn neurons in transverse spinal cord slices isolated from naïve MrgprB4-ChR2 mice (**Fig. 3A**). We observed that delivery of blue light to the spinal cord evoked post-synaptic responses in lamina I and II neurons of the superficial spinal dorsal horn. Neurons were characterized based on their action potential firing properties (**Fig. 3C)**. Firing patterns of superficial dorsal horn neurons are closely related to their neurochemical properties with tonic firing patterns being a hallmark of GABAergic inhibitory interneurons^31,32^. While postsynaptic responses were consistently observed in cell types of all firing patterns, the largest fraction of responsive-cells were comprised of tonic firing neurons (37.6%). In contrast tonic firing neurons made up only 12.5% of the non-responsive cells with the majority (62.5%) of them displaying a pattern of single firing (**Fig. 3B**). This indicates broad MrgprB4 innervation of the dorsal horn, with a potential bias towards inhibitory neuron types^33^. We then sought to investigate whether there were differences in the downstream spinal dorsal horn neurons recruited following activation of MrgprB4 lineage afferents in SNI and sham animals. Initial work has revealed that activation of MrgprB4 lineage afferents is sufficient to cause cfos expression in the superficial spinal dorsal horn (**Fig. 3D**). Ongoing work is examining the proportion of activated inhibitory and excitatory interneurons in the spinal dorsal horn following optogenetic activation of our primary afferents of interest.

**Figure 3.**
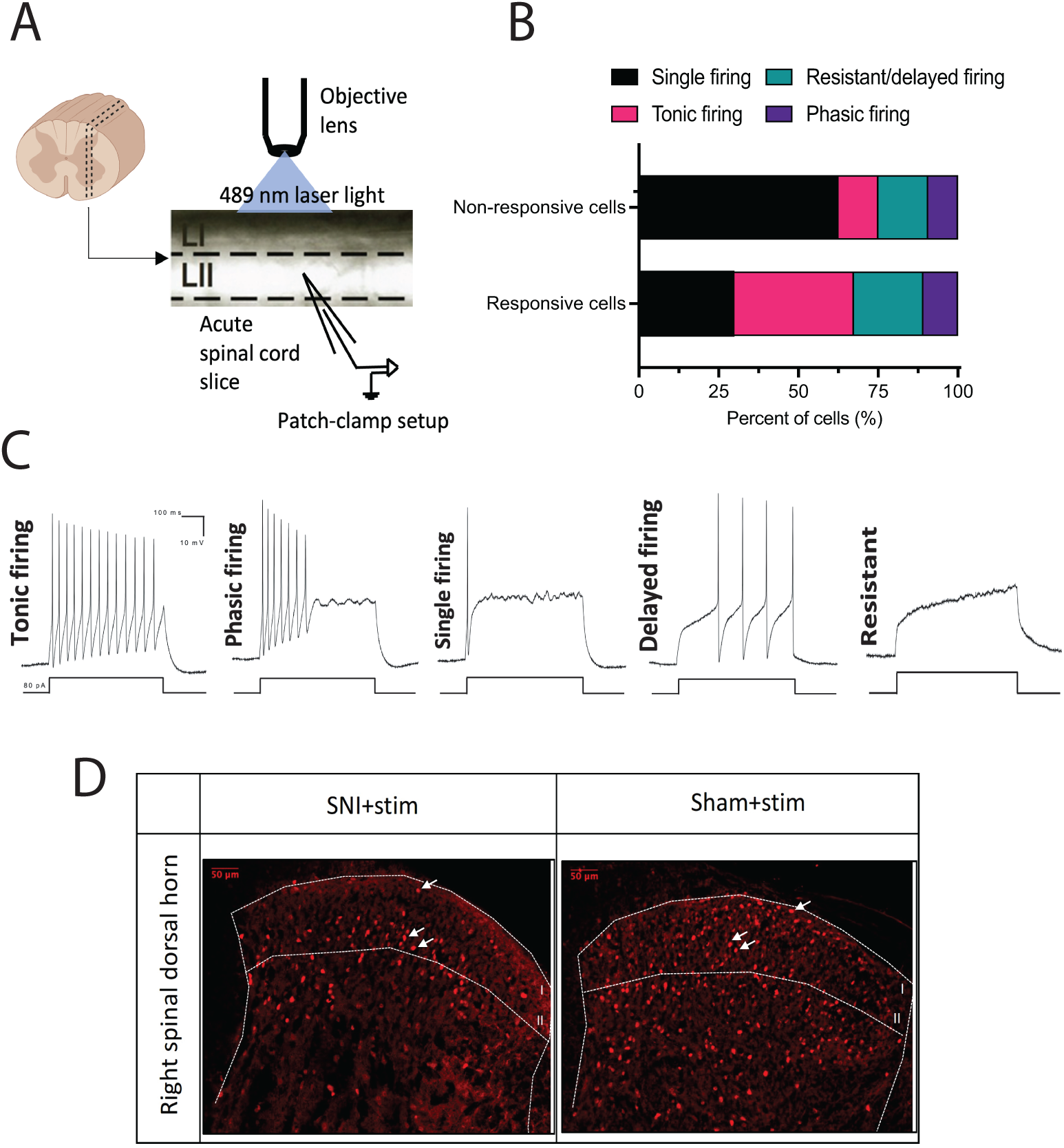
Electrophysiological characterization of the spinal dorsal horn neurons receiving input from the MrgprB4 lineage afferents. **(A)** Schematic of the recording setup used (**B**) Percent of responsive and non-responsive cells showing various firing patterns (**C**) Representative traces of the firing patterns observed. (**D**) Representative images depicting cfos expression in the right spinal dorsal horn of SNI and sham mice that received optogenetic stimulation of MrgprB4 lineage afferents.

### Differential recruitment of spinal cord projection sites in the brain following activation of MrgprB4 lineage afferents in states of nerve injury

Previous work has shown that MrgprB4 lineage afferents provide monosynaptic inputs to Gpr83+ projections neurons in the dorsal horn of the spinal cord. These neurons project to several hindbrain and midbrain targets, the most substantial input being to the lateral Parabrachial nucleus (PBN) in the hindbrain^34^. Our work showed that optogenetic activation of the MrgprB4 lineage afferents, in both nerve injured and sham mice, resulted in broad cfos expression across the lateral PBN (**Fig. 4A and B**). Intriguingly, preliminary work showed that there may be potential differences in the degree of activation in certain subdivisons of the lateral PBN between the sham and nerve injured mice(**Fig. 4A)**

**Figure 4.**
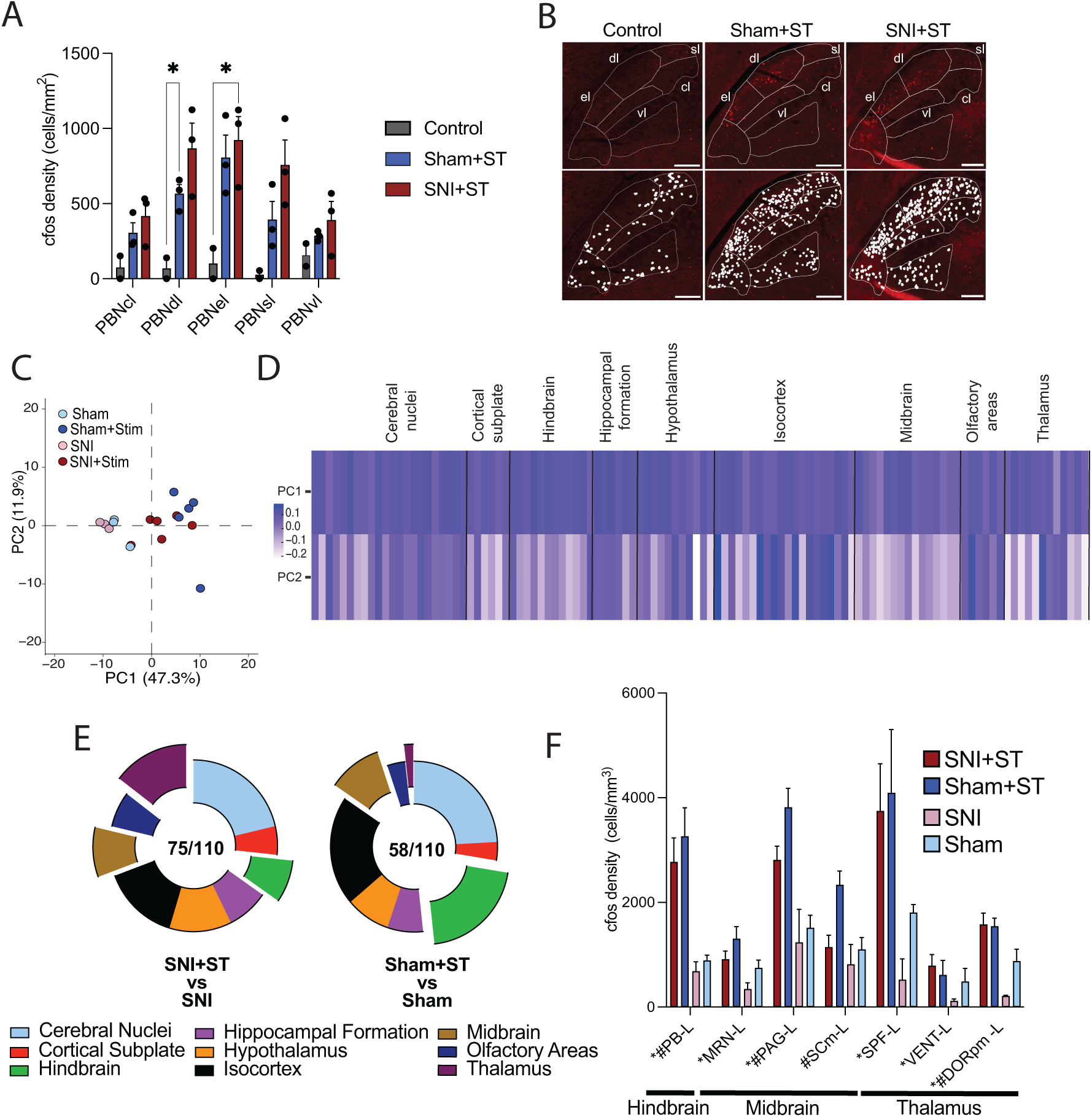
Activity patterns across the brain following activation of MrgprB4 lineage afferents in SNI and sham mice. **(A)** Quantification of cfos+ cells in various subregions of the lateral parabrachial nucleus (PBNl) in control, stimulated SNI and stimulated sham mice. RM two-way ANOVA with Tukey’s multiple comparison test (effect of brain area: *F*_(1.9, 9.4)_ = 13.5, p < 0.01; effect of experimental group: *F*_(2, 5)_ = 6.9, p < 0.05; interaction effect: *F*_(8, 20)_ = 4.6, p < 0.01; PBNdl-Control vs PBNdl-Sham+ST *p < 0.05,; PBNel-Control vs PBNel-SNI+ST *p < 0.05; 2-3 mice per group) (**B**) Representative images of the cfos staining in the PBNl with counting illustrated with solid white dots. (**C**) Scatter plot of the four conditions in the space spanned by the first two principal components of cfos density from 110 brain regions with each data point representing a mouse (**D**) Heatmap representing the loadings of individual brain regions in PC1 and PC2. (**E**) A donut plot illustrating the fraction and type of brain areas that were activated in response to stimulation in SNI and sham mice. (**F**) Bar graph depicting specific brain regions known to receive spinal cord input regarding affective touch and pain along with their results from the Mann-Whitney tests conducted on the entire dataset. * indicates significant difference between stimulated SNI compared against SNI and # indicates significant difference between stimulated sham compared against sham. Multiple Mann-Whitney tests with False discovery rate to control for multiple comparisons (n = 3-6 mice per group). el, external lateral; dl, dorsal lateral; vl, ventral lateral; cl, central lateral; sl, superior lateral; PB-L, left parabrachial nucleus; MRN-L, left midbrain reticular nucleus; SCm-L, left superior colliculus motor related; PAG-L, left periaqueductal grey; DORpm, Thalamus-polymodal association cortex related; SPF-L, left Subparafascicular nucleus; VENT-L, left ventral group of the dorsal thalamus; ST, stimulation

To assess whether the variation in appetitive coding of MrgprB4 lineage afferent activation in sham and nerve injured animals was broadly reflected across the brain we examined whole brain patterns of cfos protein in four experimental groups. These groups consisted of brains from optogenetically stimulated (MrgprB4-ChR2) and unstimulated (C57BL/6) sham and nerve injured mice that were sent to LifeCanvas Technologies for tissue clearing, cfos immunolabelling and cell counting. The acquired dataset was processed and scrubbed to include 110 regions grouped into 9 anatomical categories and all analyses were restricted to the left side which was contralateral to the side of the nerve injury.

To broadly investigate coordinated changes in activation patterns across the brain and to help inform our hypothesis testing methods we applied a principal component analysis to the raw cfos density (cells/mm^3^) values for the 110 regions from all experimental groups. The first 2 principal components comprised a majority of the variance in the dataset (>55%). A 2D projection of each mouse in the space defined by the 2 principal components revealed clear segregation between the stimulated and unstimulated groups and some partial segregation between our stimulated sham mice and our stimulated SNI mice (**Fig. 4C**). The stimulated and unstimulated groups primarily segregated along the first principal component with the stimulated SNI and stimulated sham mice partially segregating along the second principal component. To elucidate the brain-wide cfos expression pattern driving the separation of our groups, we visualized the individual loadings for all 110 brain regions (**Fig. 4D**). In general, the first PC was generated because of positive weightings across all brain areas. The PC2 loadings varied across the brain but 2 regions that displayed clusters of both positive and negative PC2 loadings were the isocortex and thalamus, implicating those regions as potential drivers of the changes in the appetitive percept following activation of MrgprB4 lineage afferents.

To statistically disentangle the exact brain areas recruited in response to optogenetic activation of MrgprB4 lineage afferents we performed multiple Mann-Whitney U tests using false discovery rate (FDR) to control for multiple comparisons (FDR = 0.1). The tests revealed no differences between the stimulated sham and stimulated SNI groups. However, several overlapping and non-overlapping areas were identified when comparing stimulated SNI against SNI and stimulated sham against sham. Broadly we observed activation in all 9 anatomical categories of brain regions with variation in the pattern of recruitment of the spinal cord projection sites as depicted by the expanded slices in the donut plot (**Fig. 4E**). A much larger fraction of thalamic sites was recruited in response to stimulation in the SNI mice compared to sham mice (11/12 compared to 1/12). Concomitantly, a larger number of hindbrain areas were recruited in response to stimulation in the sham mice compared to SNI mice (12/12 compared to 6/12). The findings regarding the thalamic sites coincide with the initial findings in the principal component analysis. To investigate which areas are uniquely activated in response to optogenetic stimulation in the sham and SNI mice we visualized the significant discoveries in our comparisons using a Venn diagram. 38 regions were found to be commonly activated in both SNI and sham mice. 37 regions were found to be uniquely activated in SNI mice and 20 regions were found to be uniquely activated in sham (SI Appendix, **Fig. 5S**). We also wanted to selectively interrogate regions that are known to receive projections from lamina I TacR1+ and/or Gpr83+ projection neurons as MrgprB4 lineage afferents are known to form monosynaptic connections with Gpr83+ neurons and potential silent polysynaptic connections with the TacR1+ neurons^34^. Consistent with our preliminary work, the parabrachial nucleus was found to be activated in response to stimulation both in the case of SNI and sham. We also saw activation across the contralateral midbrain with the midbrain reticular nucleus (MRN) recruited in case of SNI, the superior colliculus motor related (SC-m) in case of sham and periaqueductal grey (PAG) under both conditions. Within the thalamus, we saw activation of the sensory-motor cortex of the thalamus (DORsm) and polymodal association cortex of the thalamus (DORpm) in response to stimulation in SNI mice. Further subdivisions of the DORsm like the ventral group of the dorsal thalamus (VENT) and Subparafascicular nucleus (SPF) that receive input from spinal projection neurons were activated in the stimulated SNI mice. The only region of the thalamus that was activated in response to stimulation in the sham animals was the DORpm (**Fig. 4F & S4**).

## Discussion

This present study demonstrates that activation of MrgprB4 lineage primary afferents in the hindlimbs elicits appetitive touch in mice. This appetitive value was uniquely plastic and subject to change in the presence of neuropathic pain and disinhibition in the spinal cord circuits, but not acute pain induced by capsaicin. This appetitive value could be partially recovered in male nerve injured mice using gabapentin. The behavioral changes observed are also accompanied by different patterns of activity in the brain in response to optogenetic activation of MrgprB4 lineage primary afferents. This is not only true for higher order cortical structures but also subcortical and brainstem structures that receive direct input from the spinal cord.

Our immunohistochemical results along with our behavioral finding of decreased blue light preference in DIOA injected mice implicates modulation of neural signals at the spinal level as a critical player in the plasticity of the appetitive value of light touch. Disinhibition is known to unmask interconnections that allow low-threshold afferents to start signaling to neurons responsible for transmitting noxious information in the spinal cord and up to the brain^35,36^. While this is a well-established phenomenon in the realm of mechanical allodynia especially when using reflexive behavioral readouts, much less is known about how this can alter the affective coding of sensory information. Modulation of sensory signals at the level of the spinal cord can occur through spatial and temporal mechanisms where both frequency of firing and neurochemical identities of neurons can play a role in the perceptual coding of neural signals^34,37,38^.

Recent research has identified that appetitive touch is transmitted along Gpr83+ projection neurons in the spinal cord in an intensity-dependent manner where low intensity activation codes for sensory information with a positive motivational valence and high intensity activation codes for information with a negative motivational valence^34^. It is not yet known whether changes in the observed behavioral phenotype were driven by changes in the firing frequency of Gpr83+ neurons or the recruitment of the canonical nociceptive projection neurons such as the TacR1+ neurons. TacR1+ neurons have been speculated to receive silent polysynaptic input from MrgprB4 lineage neurons^34^. Disinhibition in the spinal cord circuits may unmask polysynaptic connections to TacR1+ neurons and can even increase excitatory input into Gpr83+ neurons, either of which could explain our behavioral observations. Preliminary evidence indicating differential recruitment of lateral parabrachial subnuclei, however, hints to the potential recruitment of TacR1+ neurons which innervate the parabrachial nucleus in a distinct manner compared to the Gpr83+ neurons^34^.

While the whole brain cfos dataset did not have the anatomical granularity to interrogate the subnuclei level subdivisions of the parabrachial nucleus, we still observed robust activation in the parabrachial nucleus in response to stimulation in both sham and SNI mice. A comprehensive analysis revealed several overlapping and non-overlapping regions that were activated in response to stimulation. With respect to higher order coding of sensory information in the isocortex, we found the agranular insular cortex to be commonly activated in both SNI and sham, consistent with the well-established role of the insula in affective touch processing^39–41^. The anterior cingulate cortex, prelimbic and infralimbic areas were among the isocortical regions uniquely activated in SNI, and the primary somatosensory cortex was uniquely activated in mice that had undergone sham surgery. Prelimbic and infralimbic areas have been heavily implicated in active avoidance and suppression of actions to obtain rewards in rats^42^ and the anterior cingulate cortex has long been known to play critical roles in the motivational and emotional aspects of pain^43,44^.

The aforementioned structures and the plasticity in their connections to subcortical structures has been thought to underly modulation of motivation and/or emotional valence^42,45^. Within the subcortical structures we found considerable activation across the entire thalamus and various amygdalar nuclei only in the SNI mice. These amygdalar nuclei included the central and basolateral nuclei of the amygdala whose activation has been shown to be necessary for the emergence of allodynia and depressive behaviour in animals with neuropathic pain^11,46^. While the nerve injured mice did show not show increased preference for the light paired arm in response to stimulation, they did not show heightened aversion either, using percent time as a readout. This alludes to the idea that active recruitment of specific additional structures may be driving this outwardly passive effect.

Our RTPP results in the nerve injured mice showed that while time spent in the light paired arm did not decrease with stimulation, we did observe a lower number of entries into the light paired arm in response to stimulation. A reduction in the number of entries indicates a potential subtle aversion to activation of MrgprB4 lineage neurons in states of neuropathic pain, highlighting that while MrgprB4 lineage neurons constitute a part of a circuit that transmits appetitive touch, they can also signal aversive touch under certain conditions. Emerging evidence from clinical studies have revealed the presence of anhedonia to tactile rewards such as stroking touch in chronic pain patients^12,13,47^. Chronic pain patients, particularly those with neuropathic pain, display altered evaluation of stroking touch either with respect to its perceived intensity or even its rated pleasantness^12,13^. While clinical evidence of motivational aversion to C tactile (CT) specific touch is lacking in chronic pain, research on other neurological conditions like autism show that CT specific touch can be coded as aversive^48,49^. This indicates that plasticity in skin-to-brain circuits for affective touch produces a hedonic gradient ranging from appetitive to neutral to aversive. This is especially salient considering recent preclinical and clinical findings that have shown that pathways once thought of as dedicated labelled lines for “gentle touch” are not as strict in their coding^18,22^. Redundancies in pathways for a sensation as critical as pleasurable touch and plasticity in each neuron’s ability to transmit broad modalities of cutaneous information is likely to be evolutionarily beneficial and indicates a high degree of combinatorial coding.

Notably, the appetitive value of activation of MrgprB4 lineage neurons in the RTPP was partially restored in male nerve injured mice after treatment with gabapentin. Gabapentin is a well-validated analgesic for neuropathic pain both in preclinical and clinical settings and is known to modulate both reflexive and hedonic readouts for pain. It is thought to exert its analgesic mechanism through a variety of mechanisms^50–52^. One of the primary mechanisms is its ability to block the α2δ subunit of voltage-dependent calcium channels which reduces calcium channel function and effects the spinal release of excitatory neurotransmitters^51^. It is also known to influence monoaminergic signaling in various brain regions including dopaminergic signaling in the ventral tegmental area^53^. The sexual dimorphism in gabapentin response may be driven by sex differences in the nerve injury-induced transcriptional changes in the α_2_δ subunit of voltage gated calcium channels^54^ and sex-dependent alterations in reward processing in chronic pain^55,56^. Nevertheless, these results do suggest that proper pain management approaches may ameliorate reduced appetitive processing in chronic pain and contribute to overall patient recovery.

## Materials and Methods

### Animals and housing

All experiments were approved by the University Animal Care Committee at Laval University and the Animal Ethics and Compliance Program at the University of Toronto and conducted in accordance with Canadian Council on Animal Care (CCAC) guidelines. Mice were kept on a 12:12 light:dark cycle in groups of one to four mice per cage prior to ferrule or fiber implantation, and singly housed after implantation, with food and water provided ad libitum. Adult male and/or female (2-6 months) MrgprB4-ChR2 and C57BL/6N mice were used for the experiments. Mice expressing ChR2 in MrgprB4 lineage afferents (MrgprB4-ChR2) were generated by crossing mice expressing Cre-recombinase in MrgprB4+ neurons with mice expressing a loxP-flanked STOP cassette upstream of a ChR2-EYFP fusion gene at the Rosa 26 locus (Rosa-CAG-LSL-ChR2(H134R)-EYFP-WPRE; stock number 012569, The Jackson Laboratory, Bar Harbor, ME). C57BL/6N mice were purchased from Charles River.

### Optogenetic activation of primary afferents

Depending on the needs of the experiment light delivery for optogenetic activation of the labelled primary afferents was achieved through either transdermal stimulation, an epidural implant^23^ or a vertebral implant^24^. For the central methods of light delivery, the intensity and frequency of the 470 nm laser was controlled using a LED driver (Thor Labs) and a pulse width modulator (XY-LPWM; Protosupplies, USA) or an ANY-maze Optogenetic interface (Stoelting, Co, Wood Dale, IL). In experiments with stimulation through the vertebral implant light stimulation was conducted at 3.5 mW light intensity, 2 Hz pulses and 60 sec ON 10 sec OFF. In experiments with transdermal stimulation, light stimulation was done with sustained stimulation where the lights were constantly on. In experiments with stimulation through the epidural fiber stimulation was performed at a frequency of 10 Hz. The epidural fiber implant and spinal implant surgeries were performed as previously described^23,24^. Both central methods targeted the light source to the central projections of the primary afferents originating from the hindlimbs. The mice were allowed to recover for ∼2 weeks prior to any behavioral testing.

### Electrophysiological methods

Electrophysiological recordings of miniature excitatory and inhibitory post-synaptic currents (mEPSCs and mIPSCS) and dorsal root-evoked field post-synaptic potentials (fPSPs) were made using lumbar spinal cord tissue. Adult (3−6 months old) male mice were anesthetized with urethane (2 g/kg) and perfused with ice-cold sucrose-substituted artificial cerebral spinal fluid (sucrose artificial cerebro-spinal fluid [aCSF]; contains in mM: 252 sucrose, 2.5 KCl, 1.5 CaCl_2_, 6 MgCl_2_, 10 d-glucose, bubbled with 95%:5% oxygen:CO_2_). The lumbar spinal column was removed and immersed in ice-cold sucrose aCSF after which the spinal cord was quickly removed via laminectomy. The dorsal and ventral roots were removed from the spinal cord and 300 μm-thick parasagittal slices were cut from the lumbar portion in ice-cold sucrose aCSF. After cutting, the spinal cord slices were kept in oxygenated (95% O_2_, 5% CO_2_) aCSF (in mM: 126 NaCl, 2.5 KCl, 2 MgCl_2_, 2 CaCl_2_, 1.25 NaH_2_PO_4_, 26 NaHCO_3_, 10 d-glucose) at room temperature until recording.

Slices were continuously perfused (8 ml/min) with oxygenated aCSF. Lamina II cells were visually identified using a 40X immersion objective and patched using pipettes pulled from thin-walled borosilicate glass capillary tubes. Patch pipettes had open-tip resistances of 4 to 6 MΩ when filled with an intracellular solution that contained (in mM) 145 K^+^gluconate, 5 Na^+^ gluconate, 2 KCl, 10 HEPES, 11 EGTA, 4 Mg-ATP, and 1 CaCl_2_ with an osmolarity of 300 to 320 mOsm and the pH adjusted to 7.3 with KOH. The recordings were low-pass filtered at 2 kHz and digitized at 10 kHz using a Multiclamp 700B amplifier (Molecular Devices, Sunnyvale, CA), Digidata 1322A digitizer (Molecular Devices), and pClamp 10 software (Molecular Devices). Neuronal excitability and firing properties were determined using currently clamp recording. Neurons were stimulated to action potential firing using depolarizing steps 500 ms long and increasing in 20 pA increments to a maximum of 100 pA, which was sufficient to induce action potential firing in all cells recorded. Post synaptic responses to MrgprB4-ChR2 afferent activation was determined in voltage clamp mode. ChR2-expressing afferents were activated using a 488 nm laser (Toptica iBeam) connected to the microscope. Light pulses (10 ms) were delivered to the field of view through the objective at twice the minimal light intensity threshold required to generate responses. A maximum of 20 mW was used to determine threshold and cells in which postsynaptic responses were not observed at or prior to this maximum were considered non-responsive.

### Pain models

#### Spared nerve injury

Spared nerve injury (SNI) was performed as previously described.^25^ Mice were first anesthetized with isoflurane anesthesia (3% induction and 2.5% maintenance). An initial skin incision ∼1 cm long was made longitudinally between the hip and knee joint of the right hindleg. The quadriceps femoris and biceps femoris muscles were gently mechanically separated to expose the sciatic nerve branches common peroneal, tibial, and sural. The common peroneal and tibial nerve branches were tightly ligated with a 7-0 silk suture (Ethicon, Somerville, NJ) and transected distal to the ligature. In all animals, muscles were coaxed to their original positions and skin layers were closed using interrupted 5-0 silk sutures (Ethicon). The sham surgery involved all manipulation except for the ligation and transection.

#### Intraplantar capsaicin

For the behavioral characterization of responses to transdermal stimulation following capsaicin injection a volume of 5 μL of 0.5%, wt/vol Capsaicin (dissolved in 80% saline, 10% Tween 80, and 10% ethanol) was injected subcutaneously into the plantar surface of the right hind paw. For the RTPP experiments and associated mechanical sensitivity testing a volume of 5 μL of 0.3%, wt/vol Capsaicin (dissolved in 80% saline, 10% Tween 80, and 10% ethanol) was injected subcutaneously into the plantar surface of the right hind paw. Controls for both experiments included vehicle consisting of all chemicals in the same composition without capsaicin.

#### Intrathecal DIOA

Intrathecal injections were conducted using a 30 gauge, 1/2 inch needle inserted between the L5 and L6 vertebrae using brief isoflurane anesthesia (2.5%). R-(+)-DIOA was purchased from Sigma and 2.0 mg was intrathecally administered in naive mice in a volume of 5 uL (dissolved in 10% DMSO).

### Behavioral assessment

#### Acute behavioral responses to peripheral stimulation

Acute behavioral responses to peripheral light stimulation in MrgprB4-ChR2 were measured in a binary fashion where any response or movement occurring during or immediately after light exposure was rated as a 1 and a lack of response was rated as 0. This scoring was performed at increasing discrete light intensities.

#### Real time place preference

To study the behavioral output of activation of the MrgprB4 lineage primary afferents we use a real time place preference (RTPP) assay with light delivery through the spinal implant. The assay is a V shaped maze with 2 different visual contexts in each arm. The implanted and tethered mouse is placed in the assay and light delivery is controlled through the ANY-maze video tracking software. Light is activated (3.5 mW, 2 Hz, 60 s ON 10s OFF) when the mouse enters the light paired arm and deactivated when the mouse exits the light paired arm. Baseline preference of arms without any light stimulation is also initially recorded. Since the mice showed different baseline preferences for each arm, we used a biased experimental design. We perform light stimulation in the initially non-preferred (INP) arm to assess whether we can increase the preference for that arm by associating it with activation of MrgprB4 lineage primary afferents.

A variation of the real time place preference (RTPP) assay was also created for transdermal stimulation. It consisted of a rectangular arena with an LED-illuminated floor. Half of the arena was illuminated with blue light and the other half was illuminated with orange light. This color differential was achieved by placing blue (400-460 nm) and orange filters (580-740 nm) over the LEDs. Mice were tracked for a total of 20 minutes; 10 minutes before and 10 minutes during illumination, to assess preference for the illuminated blue or orange zone.

#### Mechanical sensitivity – Von Frey

Mechanosensitivity was assessed using von Frey filaments. Mice were placed in a small chamber with a grid floor and allowed to habituate for 30 minutes. The affected hind paws were tested using the SUDO method^57^ and the paw withdrawal threshold was reported in pressure (g/mm2)^58^.

#### Thermal sensitivity – Hargreaves

Thermal sensitivity was assessed using a Hargreaves thermal sensitivity apparatus (IITC Life Sciences). Mice were placed on a 3/16th-inch thick glass floor warmed to 29℃ within small (8 × 8 cm) Plexiglas cubicles, and a focused high-intensity projector lamp beam was shone from below onto the mid-plantar surface of the hind paw.

### Immunofluorescence – In house

Brains, spinal cords and lumbar (L5-L6) dorsal root ganglions (DRG) of male and female mice were harvested immediately after transcardial perfusion with PBS followed by 4% PFA in PBS. The spinal cords and DRGs were post fixed in 4% PFA for 4 hours at 4 °C and brains were fixed overnight at 4 °C. The tissue was washed in 5 times in PBS and then subsequently placed in 30% sucrose for 48 hours. The tissue was then flash frozen in OCT using dry ice.

Brains and spinal cords were sectioned at 30 um directly onto superfrost slides. DRGs were sectioned at 16 um directly onto superfrost slides. All slides were placed in the −20 freeze. The day of the staining the slides were removed from the freezer and placed in a humidified chamber to come to room temperature. The slides were washed 3x 10 min with PBST followed by the application of the primary antibody in PBST with 5% horse serum. The primary antibodies were left overnight at room temperature. After 3x 10 min PBST washes the secondary antibodies in PBST were applied for 1-2 hours at room temperature. Slides were washed 3x 10 min in PBST, followed by application of DAPI (1:500 in PBS) for 10 min and the 3x 5 min PBS washes. Slides were then mounted with a coverslip and mounting medium. The primary antibodies used included Mouse Anti-NeuN (1:1000; MAB377, Chemicon), Chicken Anti-GFP (1:1000; ab13970, Abcam), Rabbit Anti-cfos (1:1000; ab190289, Abcam), Rabbit Anti-ATF3 (1:500; ab207434, Abcam). The secondary antibodies used included Alexa-Fluor 647 anti-Rabbit (1:500; ab150075, Abcam), Alexa-Fluor 488 anti-Chicken (1:500; 144438, Jackson Immuno Research) and Alexa-Fluor 488 anti-Mouse (1:500; 615-545-214, Jackson Immuno Research).

### Whole mouse brain processing, staining, and imaging – LifeCanvas technologies

SHIELD-based whole-brain clearing and labeling was employed for mapping cFos activity in sham and SNI male mice that either received optogenetic stimulation through a spinal implant (MrgprB4-ChR2) or did not have a spinal implant and received no stimulation (C56BL/6). Time post-surgery, handling and housing conditions were kept consistent across all 4 groups of mice.

Perfusion and dissection was performed 50 minutes after a 10 minute optogenetic stimulation session. Brains were harvested immediately after transcardial perfusion with PBS + heparin followed by perfusion with 4% PFA in PBS. The brains were fixed in 4% PFA at 4°C for 24 hours with gentle shaking. Samples were washed 5 times in PBS and then shipped to LifeCanvas technologies in PBS + 0.02% of sodium azide for further processing.

Samples were cleared for 7 days with the Clear+ delipidation buffer. The samples were batch labelled with SmartBatch+ using 10 ug Mouse NeuN and 3.5 ug Rabbit cFos antibody per brain. The brains were imaged with SmartSPIM at 4 um z-step and 1.8 um xy pixel size.

Sample images were tile corrected, destriped and registered to the Allen Brain Atlas (Allen Institute: https://portal.brain-map.org/) using an automated process. A NeuN channel for each brain was registered to 8-20 atlas-aligned reference samples, using successive rigid, affine, and b-spline warping algorithms (SimpleElastix: https://simpleelastix.github.io/). An average alignment to the atlas was generated across all intermediate reference sample alignments to serve as the final atlas alignment value for the sample. Automated cell detection was performed using a custom convolutional neural network through the Tensorflow python package. The cell detection was performed by two networks in sequence. First, a fully-convolutional detection network (https://arxiv.org/abs/1605.06211v1) based on a U-Net architecture (https://arxiv.org/abs/1505.04597v1) was used to find possible cell-positive locations. Then a convolutional network using a ResNet architecture (https://arxiv.org/abs/1512.03385v1) was used to classify each location as positive or negative. Using the atlas registration, each cell location was projected onto the Allen Brain Atlas to count the number of cells for each atlas-defined region.

### Statistical methods

In the real time place preference experiments mice displaying extreme preferences for either arm or side (>95% of the time) were excluded from the analysis. The experimenters were blinded to experimental groups during testing, video analysis and image analysis. Statistical analyses were performed using GraphPad Prism v10 and RStudio. In all figures, results are expressed as the means ± SEM or as the mean and spread of the data points. Behavioral data was analyzed using paired t-tests, log-rank tests, repeated measures one-way or two-way ANOVA with Sidak multiple comparison tests. Immunohistochemical data from the DRG slices was analyzed using unpaired t-tests. Immunohistochemical data from slices were analyzed using Repeated measures two-way ANOVA with Tukey’s multiple comparison test. Immunohistochemical data from whole brain cfos was analyzed with principal component analysis using Rstudio and with multiple Mann Whitney U tests (with FDR = 0.1) in GraphPad Prism.

