## Supplemental Figures for "Chronic pain mediated changes in the appetitive value of affective gentle touch in mice"

**Supplemental information (SI appendix)**

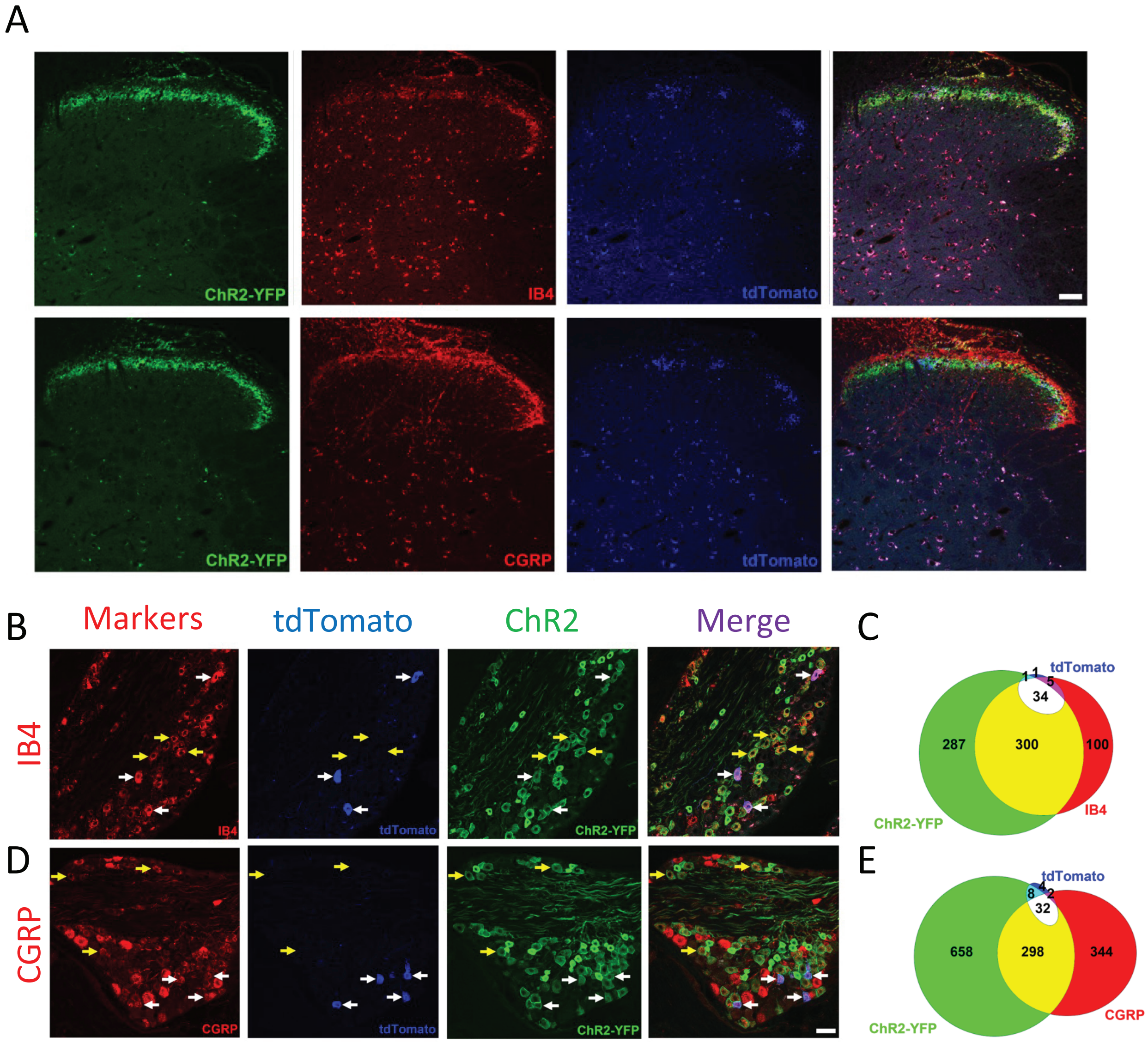


**Figure S1. Characterizing expression of ChR2-YFP in MrgprB4 lineage neurons.**
**(A)** Innervation pattern of the MrgprB4 lineage neurons in the superficial dorsal horn of the spinal cord. **(B and C)** Simultaneous immunostaining of IB4(non-peptidergic afferents), tdTomato and ChR2-YFP and the degree of overlap between the 3 groups in the dorsal root ganglions (DRG) of MrgprB4-ChR2 mice. **(D and E)** Simultaneous immunostaining of CGRP(peptidergic afferents), tdTomato and ChR2-YFP and the degree of overlap between the 3 groups.
Yellow arrows indicate overlap of ChR2 and the marker and white arrows indicate ChR2, tdTomato and marker overlap.

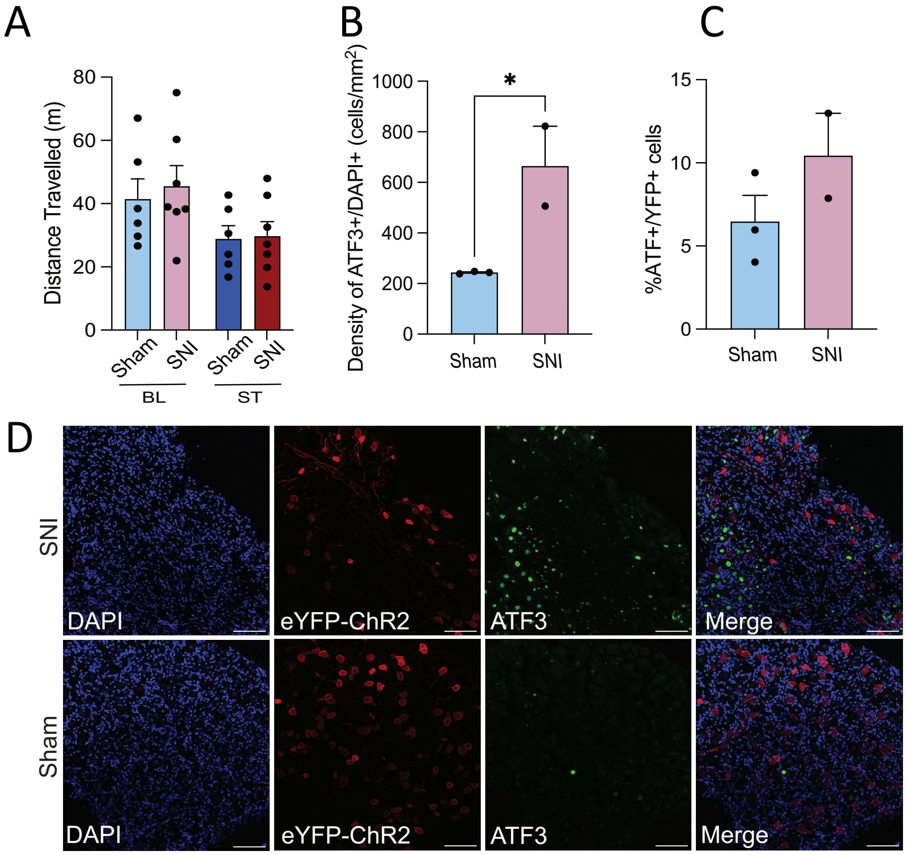


**Figure S2. Characterizing behavioral and molecular features of the SNI mice, related to Figure 2.** **(A)** No difference in distance travelled between SNI and sham mice both during baseline and during the stimulation trial. RM two-way ANOVA with Sidak multiple comparison test (effect of trial: $F_{(1, 11)} = 8.2$, $p < 0.05$; effect of surgery: $F_{(1, 11)} = 0.2$, $p > 0.5$; interaction effect: $F_{(1, 11)} = 0.1$, $p > 0.5$; n = 6-7 per group) **(B)** Quantification of the density of ATF3+ cells that are also DAPI+, depicting higher expression of a nerve injury marker, ATF3, in SNI mice compared to sham. **(C)** Quantification of the percent of YFP+ cells that are also ATF3+. Unpaired t-test ($p > 0.05$; n = 2-3 mice per group). (**D**) Immunofluorescence images displaying DAPI, eYFP-ChR2 and ATF3 expression in both groups. Scale bars represent 100 um.

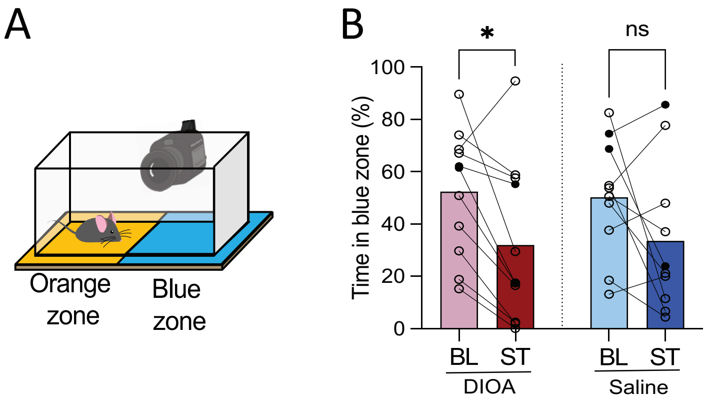


**Figure S3. Decreased preference for the blue light zone in mice with pharmacological disinhibition. (A)** Illustration of the peripheral light-based real-time place preference assay used. (**B**) Decreased preference for blue side during stimulation compared to baseline for the DIOA injected mice but not the saline injected mice. RM two-way ANOVA with Sidak multiple comparison test (effect of trial**:** $F_{(1, 19)} = 10.2$, $p < 0.01$; effect of injection**:** $F_{(1, 19)} = 0.00007$, $p > 0.05$; interaction effect**:** $F_{(1, 19)} = 0.1$, $p > 0.05$; Saline-BL vs Saline-ST $p > 0.05$, DIOA-BL vs DIOA-ST *$p < 0.05$; n = 10-11 mice per group).

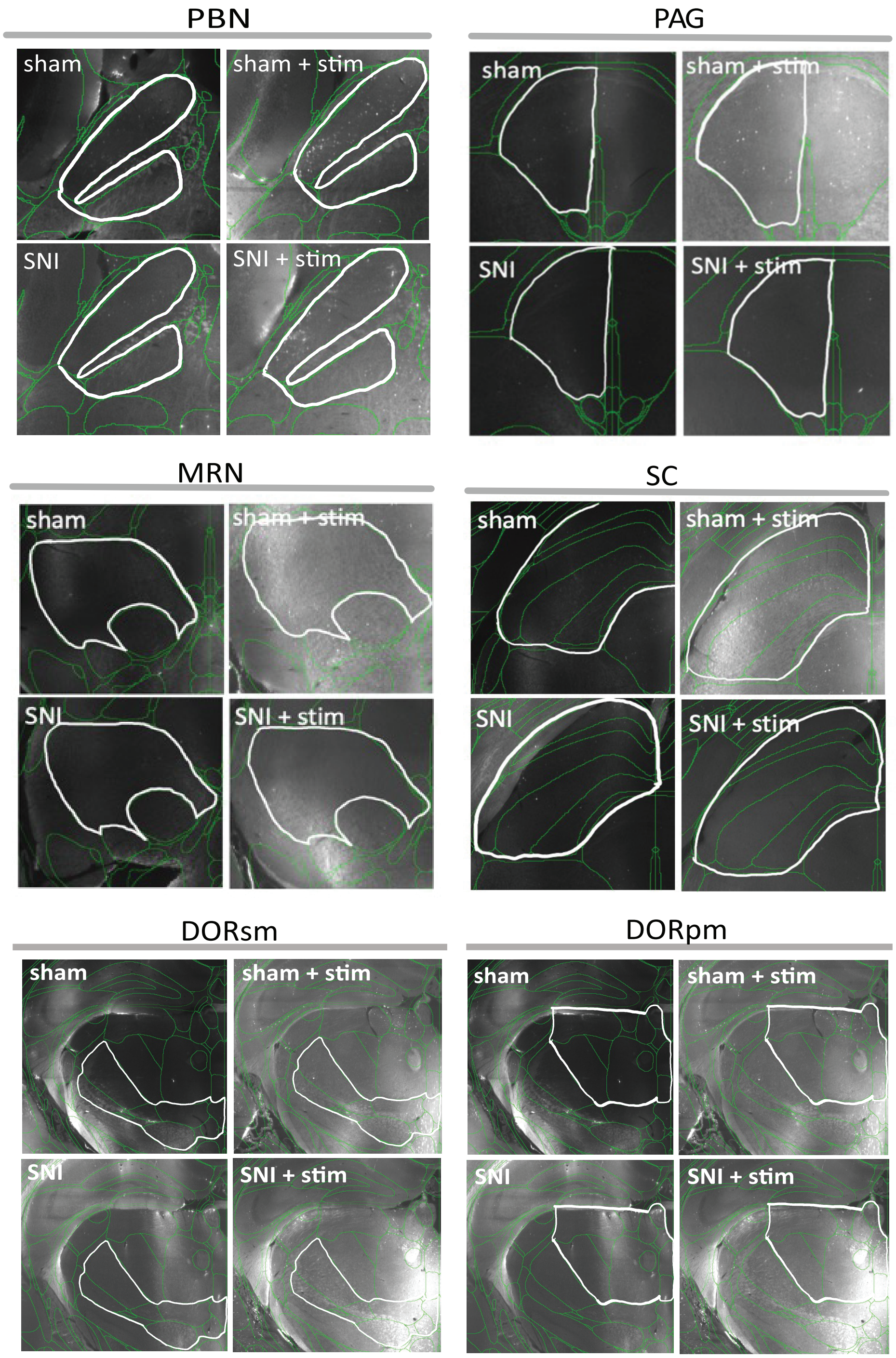


**Figure 4S. Representative images from whole brain cfos following stimulation in SNI and sham animals, related to Figure 4.** Images depict cfos expression in the contralateral PBN of the hindbrain, PAG, MRN & SC of the midbrain and the DORsm & DORpm of the thalamus. PBN, left parabrachial nucleus; MRN, left midbrain reticular nucleus; SC, left superior colliculus; PAG, left periaqueductal grey; DORpm, Thalamus – polymodal association cortex related; DORsm, Thalamus – sensory motor cortex related

**Chronic pain mediated changes in the appetitive value of affective touch in mice**
Zain et al 2023

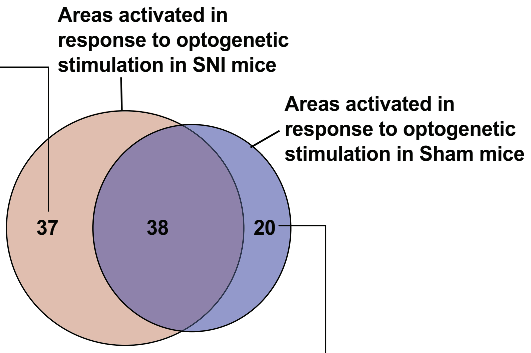


**Cerebral Nuclei**
1. Central amygdalar nucleus
2. Intercalated amygdalar nucleus
3. Bed nucleus of the anterior commissure
4. Magnocellular nucleus

**Cortical Subplate**
1. Basolateral amygdalar nucleus
2. Basomedial amygdalar nucleus
3. Posterior amygdalar nucleus

**Hippocampal Formation**
1. Field CA2
2. Parasubiculum

**Hypothalamus**
1. Hypothalamic lateral zone
2. Paraventricular hypothalamic nucleus - descending division
3. Zona incerta
4. Paraventricular hypothalamic nucleus
5. Periventricular zone
6. Subthalamic nucleus

**Isocortex**
1. Ectorhinal area
2. Infralimbic area
3. Perirhinal area
4. Prelimbic area
5. Anterior cingulate area
6. Orbital area medial part

**Midbrain**
1. Red nucleus
2. Midbrain reticular nucleus
3. Ventral tegmental area

**Olfactory areas**
1. Dorsal peduncular area
2. Piriform-amygdalar area
3. Cortical amygdalar area

**Thalamus**
1. Thalamus sensory-motor cortex related
2. Geniculate group of the ventral thalamus
3. Lateral group of the dorsal thalamus
4. Lateral habenula
5. Medial group of the dorsal thalamus
6. Midline group of the dorsal thalamus
7. Reticular nucleus of the thalamus
8. Subparafascicular nucleus
9. Ventral group of the dorsal thalamus
10. Anterior group of the dorsal thalamus

**Cerebral Nuclei**
1. Bed nucleus of the accessory olfactory tract
2. Medial septal complex

**Cortical Subplate**
1. Endopiriform nucleus

**Hindbrain**
1. Intermediate reticular nucleus
2. Laterodorsal tegmental nucleus
3. Nucleus incertus
4. Nucleus of the solitary tract
5. Nucleus raphe pontis
6. Sublaterodorsal nucleus

**Hypothalamus**
1. Arcuate hypothalamic nucleus
2. Ventromedial hypothalamic nucleus

**Isocortex**
1. Gustatory areas
2. Orbital area
3. Orbital area lateral part
4. Primary motor area
5. Primary somatosensory area
6. Secondary motor area
7. Somatosensory areas

**Midbrain**
1. Superior colliculus motor related
2. Superior colliculus sensory related

**Figure 5S. Non-overlapping brain areas activated in response to optogenetic stimulation of MrgprB4 lineage afferents in sham and SNI mice.**
